# In-depth specificity profiling of Pro-Pro endopeptidases (PPEPs) using combinatorial synthetic peptide libraries

**DOI:** 10.1101/2022.11.15.516584

**Authors:** Bart Claushuis, Robert A. Cordfunke, Arnoud H. de Ru, Annemarie Otte, Hans C. van Leeuwen, Oleg I. Klychnikov, Peter A. van Veelen, Jeroen Corver, Jan W. Drijfhout, Paul J. Hensbergen

## Abstract

Proteases comprise the class of enzymes that catalyze the hydrolysis of peptide bonds, thereby playing a pivotal role in many aspects of life. The amino acids surrounding the scissile bond determine the susceptibility towards protease-mediated hydrolysis. A detailed understanding of the cleavage specificity of a protease can lead to the identification of its endogenous substrates, while it is also essential for the design of inhibitors. We developed a new method which combines the high diversity of a combinatorial synthetic peptide library with the sensitivity and detection power of mass spectrometry to determine protease cleavage specificity. We applied this method to study a group of bacterial metalloproteases that have the unique specificity to cleave between two prolines, i.e. Pro-Pro endopeptidases (PPEPs). We not only confirmed the prime-side specificity of PPEP-1 and PPEP-2, but also revealed some new unexpected peptide substrates. Moreover, we have characterized a new PPEP (PPEP-3) which has a prime-side specificity that is very different from that of the other two PPEPs. Importantly, the approach that we present in this study is generic and can be extended to investigate the specificity of other proteases.

## Introduction

Proteases comprise the class of enzymes that catalyze the hydrolysis of peptide bonds between amino acids in a polypeptide chain. Through cleavage of their substrates, proteases play a pivotal role in many aspects of life, ranging from viral polyprotein processing^1^ to a wide range of human physiological and cellular processes, e.g. hemostasis, apoptosis and immune responses^2–4^. Substrate specificity of a protease is primarily determined by the interaction between the substrate and the protease within the active site. The amino acids at positions N-terminal (P1-P4) and C-terminal (P1’-P4’) to the scissile bond (P1-P1’) generally determine the susceptibility of the polypeptide towards protease-mediated hydrolysis, but the length and composition of the peptide cleavage consensus depends on the protease. Some proteases are very specific, i.e. they have only one or a very limited number of substrates, while others are more promiscuous, such as digestive enzymes.

Uncovering the endogenous substrate(s) is usually a key step towards dissecting the biological role of a protease. However, it is not easy to identify protease substrates without prior knowledge, e.g. without a clear phenotype in a protease knockout or lack of information from homologs in other species. Information about the cleavage specificity of a protease can also aid in the identification of endogenous substrates. Moreover, such information is pivotal for inhibitor design or the development of diagnostic biomarker assays^5–7^. Hence, numerous methods for protease specificity profiling have been developed^6,8^. Some of these make use of the high diversity of libraries, either by phage display technologies^9,10^ or as a large collection of synthetic peptides.

As a read-out for cleavage of substrates in peptide libraries, fluorescence detection^5,11^, Edman degradation^12,13^ and mass spectrometry (MS)^14–17^ have been used. The latter is especially attractive because it determines the signature proteolytic event in a highly specific manner, i.e. information on the amino acid(s) surrounding the scissile bond is obtained. Moreover, recent advances in the speed and sensitivity of the current MS instruments enable robust and versatile analyses. Obviously, the overall diversity of peptides within the library is an important parameter, not only to assure a potential substrate is represented and therefore protease activity can be detected, but also to gain knowledge about the specificity of the protease for certain positions surrounding the scissile bond.

Combinatorial synthetic peptide libraries seek to achieve the full range of possible substrates by randomizing the positions surrounding the cleavage site^11,18^. To our knowledge, MS has hitherto not been used to detect product formation following incubation of such a library with a protease of interest. Obviously, the complexity of combinatorial libraries grows rapidly when increasing the number of positions that are to be randomized. Therefore, minimizing the variety of amino acids at certain positions in the peptides and enrichment of product peptides are preferred. Partial randomization would be an optimal research strategy for the group of bacterial proteases, which we have previously identified, that have the unique specificity to cleave a peptide bond between two prolines, i.e. Pro-Pro endopeptidases (PPEPs).

The first two members of this family of proteases that were identified, PPEP-1 from the human pathogen *Clostridioides difficile*^19,20^ and PPEP-2 from *Paenibacillus alvei*^21^, are secreted and cleave cell surface proteins involved in bacterial adhesion. Initially, the specificity of PPEP-1 was determined based on a small synthetic peptide library that was designed based on the identification of a sub-optimal cleavage site in a human protein^22^. Following the elucidation of the endogenous PPEP-1 substrates, in which a total of 13 cleavage sites were found, a cleavage motif could be determined (Fig. 1a). For PPEP-2, the endogenous cleavage site (Fig. 1a) was experimentally determined following an *in silico* prediction of the substrate. This prediction was based on a similar genomic organization of the PPEP gene and its substrate in both *C. difficile* and *P. alvei,* i.e. they are adjacent genes (Fig. 1b). Based on a bioinformatic analysis, we recently observed PPEP homologs in a wide variety of species^23^, for example in *Geobacillus thermodenitrificans* (PPEP-3, Fig. 1). The modeled structure of PPEP-3 shows a high degree of similarity with the crystal structures of PPEP-1 and PPEP-2 (Fig. 1c). However, none of the genes adjacent to *ppep-3* encode a protein which contains a PPEP consensus cleavage motif (XXPPXP, Figs. 1a,b), hampering the formulation of a testable hypothesis about its substrate(s). Hence, to gain insight in the activity and specificity of hitherto uncharacterized putative PPEPs, a general method to profile their specificity is needed.

**Fig 1.**
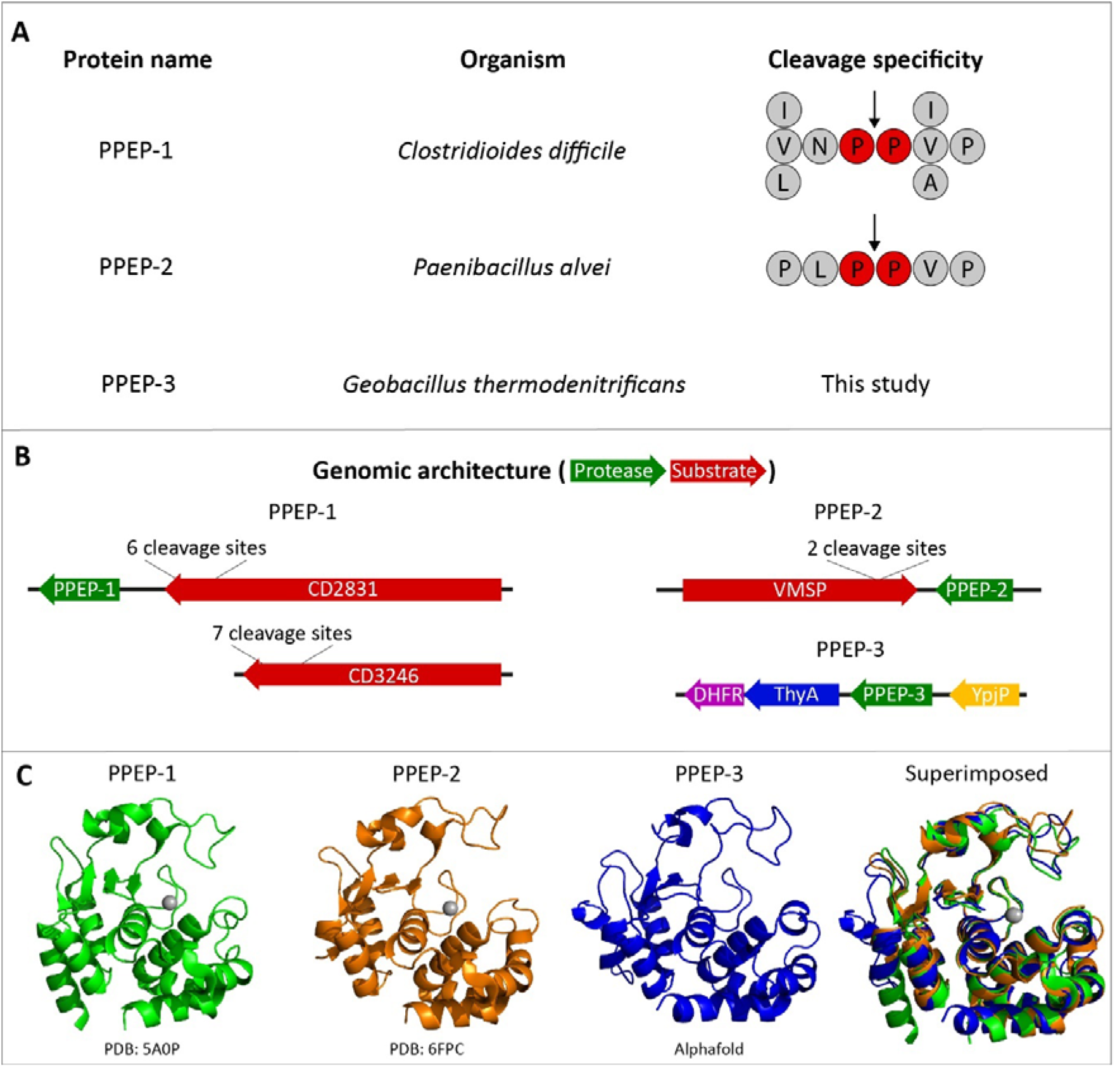
Overview of the PPEPs used in this study. **a** The three PPEPs that are used in this study and their respective origins and substrate specificity. For PPEP-1 and 2 the cleavage specificity is based on the endogenous substrates. For PPEP-3, no substrates have been described yet. **b** The genomic architecture of the PPEPs and their substrates. For PPEP-1, the gene encoding the substrate CD2831 is adjacent to PPEP-1. The gene encoding the second substrate (CD3246) is positioned elsewhere on the genome. The genes for PPEP-2 and its substrate VMSP are also located adjacent to each other. For PPEP-3, no adjacent genes contain the consensus PPEP cleavage motif (i.e. PPXP). **c** Crystal (PPEP-1 and PPEP-2) and predicted (PPEP-3) structures^20,21^. PPEP-3 structure was predicted using the Alphafold algorithm^24^.

Therefore, the aim of the current study was to develop a method, which combines the high diversity of a combinatorial peptide library with the sensitivity and specificity of MS detection, to study the activity and specificity of PPEPs. Testing the method with PPEP-1 and PPEP-2 showed results that were in good agreement with previous data, while also some unexpected peptide substrates were observed. Importantly, the new method clearly established PPEP-3 as a genuine PPEP, but also showed that it has a markedly different prime-side specificity compared to PPEP-1 and PPEP-2.

## Results

### Combinatorial peptide library design and experimental setup

Since PPEPs are defined by their ability to hydrolyze Pro-Pro bonds, and substrate specificity is further determined by positions P3-P3’ surrounding the scissile bond^20,22,25^, we constructed a combinatorial peptide library containing a XXPPXX motif. In this motif, the X positions represent any amino acid residue (with the exception of cysteine), while the core proline (P) residues, corresponding to the P1-P1’ positions, are fixed (Fig. 2).

**Fig. 2.**
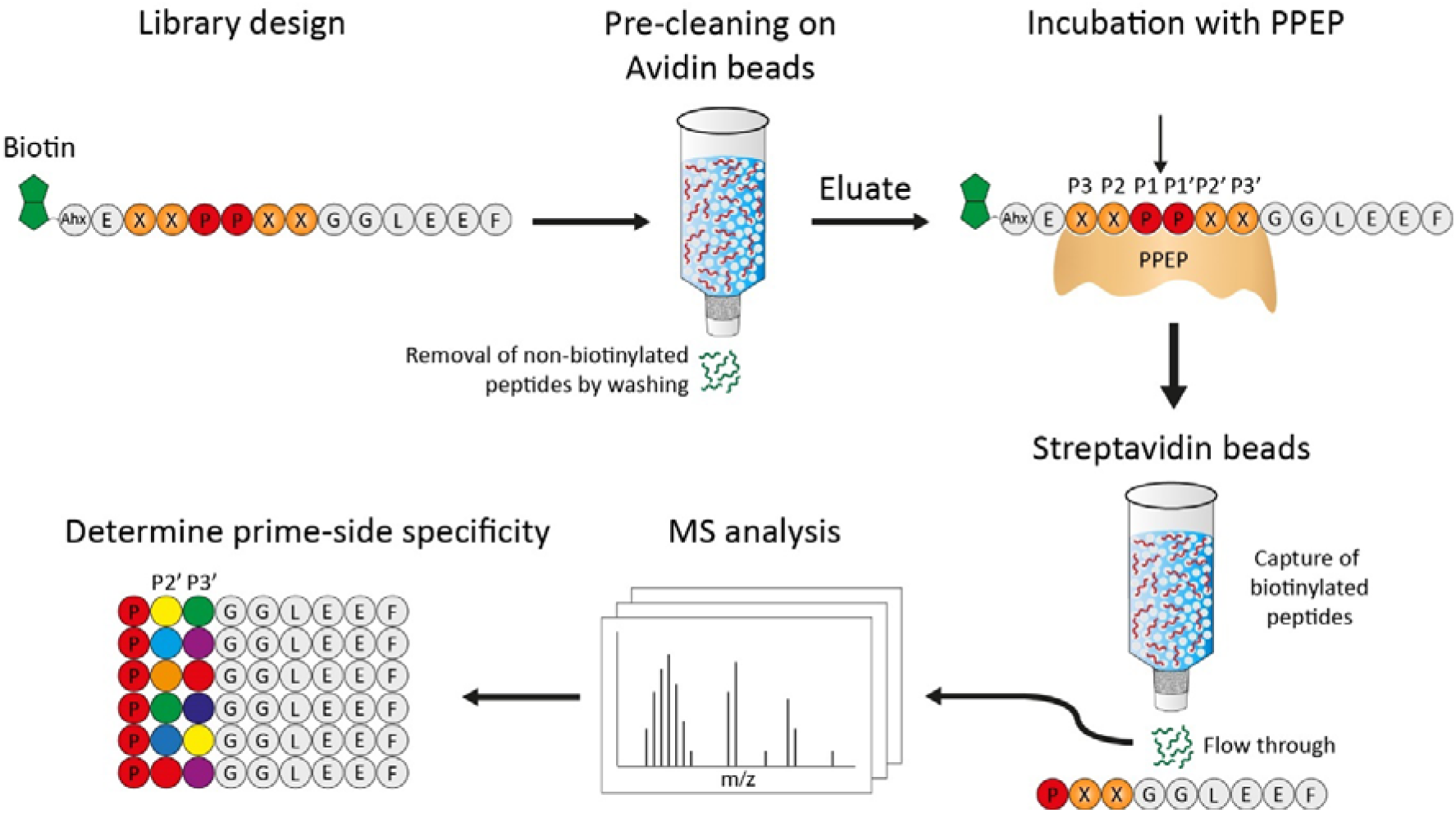
Design of the synthetic combinatorial peptide library and workflow to determine the activity and prime-side specificity of a Pro-Pro endopeptidase (PPEP). The library was designed to contain an XXPPXX motif, X representing any residue (X≠Cys). First, the library was pre-cleaned on avidin beads to remove non-biotinylated peptides. Then, the library was incubated with a PPEP. The scissile bond is indicated by the arrow. Following this, biotinylated peptides (non-cleaved peptides and N-terminal product peptides) were captured on a streptavidin column. The flow-through, containing non-biotinylated C-terminal product peptides (PXXGGLEEF) were then analyzed by LC-MS/MS, after which the prime-side specificity could be determined. Ahx: 1-aminohexanoic acid.

In order to analyze product peptides after incubation of the library with a PPEP, the core sequence (XXPPXX) was modified in two ways. First, a six amino acid tail consisting of Gly-Gly-Leu-Glu-Glu-Phe (GGLEEF) was added at the C-terminus (Fig. 2). This sequence was chosen because PPEP cleavage between the two prolines would then provide retention of the C-terminal product peptides (PXXGGLEEF) on a C18-column. Moreover, the fragmentation pattern of such a peptide (PYVGGLEEF) that we observed in a previous study provided good sequence coverage of the N-terminal region (Supplementary Fig. 1). Second, a biotin was attached to the N-terminus of each peptide, connected to the rest of the peptide by a small linker (Ahx-Glu, Ahx=1-aminohexanoic acid, Fig. 2). This allows for the enrichment of C-terminal product peptides by removal of biotinylated peptide molecules, i.e. non-cleaved peptides and N-terminal product peptides, using streptavidin beads.

Synthesis of the library was performed using the one-bead one-compound (OBOC) method^26^ in order to achieve equimolar amounts of each unique peptide. Initially, we synthesized 19 sub-libraries for which the amino acid at the X corresponding to the P3 position (the first X in the sequence XXPPXX) was known. Each of these sub-libraries contains 6859 peptides (19×19×19). Since the process of linking biotin to the N-terminus is not 100% efficient, nonbiotinylated peptides were also present. To remove these unwanted peptides prior to incubation with a PPEP, the library was pre-cleaned on an avidin column (Fig. 2). The biotinylated peptide library that was obtained after elution from the avidin column was then incubated with a PPEP and subsequently depleted for biotinylated peptides using streptavidin. C-terminal, non-biotinylated, product peptides (PXXGGLEEF) were collected in the flow-through and analyzed by mass spectrometry. Following data analysis, the amino acids at the P2’ and P3’ positions were determined (Fig. 2).

### Incubation of PPEP-1 with two sub-libraries confirms the preference of PPEP-1 for valine over lysine at the P3 position

In our previous studies, we showed a preference of PPEP-1 for a Val as compared to a Lys at the P3-position^25^. Hence, to test the feasibility of our approach, two sub-libraries with either a Val or Lys at this position were incubated with PPEP-1. The formation of products due to proteolysis of substrate peptides present in the library was assessed using MALDI-FT-ICR MS (Fig. 3). As expected, product peptides were clearly visible when using the P3=Val library (Fig. 3, upper panel), while these were not observed when the P3=Lys library was used instead (Fig. 3, lower panel).

**Fig. 3.**
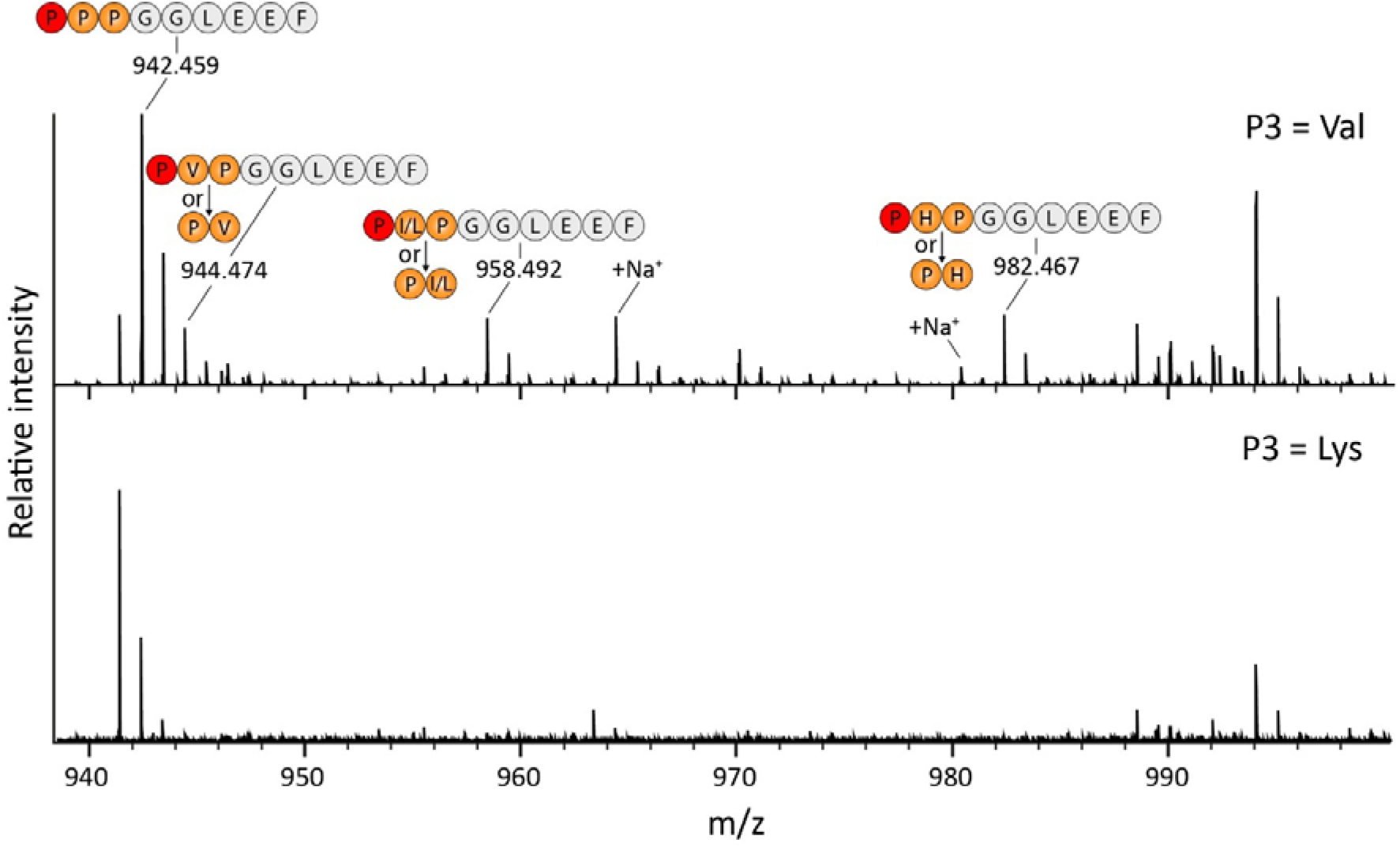
MALDI-FT-ICR MS analysis of PPEP-1 product peptides using two different combinatorial sub-libraries. The P3=Val and P3=Lys sub-libraries were incubated with PPEP-1 for 3 h. Following depletion of biotinylated peptides, non-biotinylated product peptides (PXXGGLEEF) were analyzed using MALDI-FT-ICR MS. The two indicated sodiated species are from the PPPGGLEEG and P(I/L)PGGLEEF/(PP(I/L)GGLEEF peptides, respectively.

Although no fragmentation was performed, we could assign several product peptides when using the P3=Val library based on the accurate mass and our current understanding of the specificity of PPEP-1 (Fig. 1)^22,25^. The highest signal was observed for the PPPGGLEEF peptide (*m/z* 942.459, [M+H]^+^). Although three prolines at P1’-P3’ are not found in the endogenous substrates (Fig. 1), it had been demonstrated that PPEP-1 prefers all prolines at these positions^22^. In addition, a peptide matching with the product peptide PIPGGLEEF was observed, although based on the MALDI-FT-ICR MS analysis alone we cannot exclude the possibility that it corresponds to PPIGGLEEF, nor that it might contain a leucine instead of an isoleucine at the site corresponding to the P2’/P3’ position. We also observed a peptide corresponding to PVPGGLEEF (or PPVGGLEEF). Even though the signal for this peptide partially overlapped with the second isotope peak of the PPPGGLEEF peptide (theoretical *m/z* value: 944.462, [M+H]^+^), a separate peak for the signal at *m/z* 944.474 ([M+H]^+^) was clearly visible. Lastly, a peptide was observed corresponding to either PHPGGLEEF or PPHGGLEEF even though it was hitherto unknown that PPEP-1 allows for a histidine at the P2’ or P3’ position.

Overall, the above results with the two combinatorial sub-libraries demonstrated the applicability of our approach to detect PPEP activity and study its preference for amino acids surrounding the scissile Pro-Pro bond.

### PPEP-1, PPEP-2 and PPEP-3 display distinct substrate specificity after incubation with the full combinatorial peptide library

Following the successful tests of the method with the two sub-libraries and PPEP-1, we applied our method with the full combinatorial peptide library (a mix of all 19 sub-libraries, containing 130,321 peptides) to determine the prime-side substrate specificity of PPEP-1, PPEP-2 and PPEP-3. In order to increase the sensitivity and include fragmentation of the product peptides, we analyzed the samples with LC-MS/MS. A non-treated sample was included as a control.

Initially, we analyzed the results by standard database searching against an in-house generated database (see Methods for details). For PPEP-1 and PPEP-2 treated samples, the peptides with the highest intensities represented the expected PXXGGLEEF product peptides (Supplementary Table 1). Moreover, an enrichment for prolines at the P2’ and/or P3’positions was observed (Supplementary Table 1), in line with what was expected based on the specificity of PPEP-1 and PPEP-2 (Fig. 1). For the PPEP-3 treated sample, the most highly abundant peptide was PPPGGLEEF. Hence, this clearly demonstrated that also PPEP-3 is an authentic PPEP. In addition, other 9-mer PXXGGLEEF product peptides were present among the most abundant peptides in the PPEP-3 treated sample (Supplementary Table 1).

The results from the database search showed ambiguity in the position of the proline at the P2’/P3’ position as assigned by the search algorithm (i.e. PXPGGLEEF or PPXGGLEEF). Also, several MS/MS spectra were matched with sequences that did not match with the expected 9-mer PXXGGLEEF sequence. For example, some MS/MS spectra were assigned to the 8-mer sequence KYGGLEEF. However, we argue that these represent wrong annotations due to the fact that the mass and elution time of this peptide is exactly the same as the PPPGGLEEF peptide, (one of) the highest product peptides observed for all three PPEPs (Supplementary Table 1). Furthermore, in all cases that an isoleucine or leucine was present at the P2’ or P3’ position, obviously no distinction could be made by the search algorithm.

To substantiate our results, we combined manual inspection of the MS/MS spectra with additional LC-MS/MS analyses of a set of synthetic peptides. Fortunately, fragmentation spectra of PXPGGLEEF and PPXGGLEEF peptides showed clear differences (Supplementary Fig. 2). Moreover, isomeric peptides could often be separated on a C18 column (e.g. PHPGGLEEF vs PPHGGLEEF and PLPGGLEEF vs PIPGGLEEF, Supplementary Fig. 3). Based on these additional analyses, we could refine the results from the database search and accurately assign the identity and abundance of the individual product peptides, thereby allowing us to determine the prime-side specificity for the three PPEPs (Fig. 4). Although we observed some longer peptides (Supplementary Table 1), e.g. the 10-mer peptide PPPPGGLEEF) we focused our attention to the top 10 most highly abundant 9-mer product peptides. Of note, since the proline at the P1’ was fixed (Fig. 2), no variation is observed at this position in Fig. 4.

**Fig. 4.**
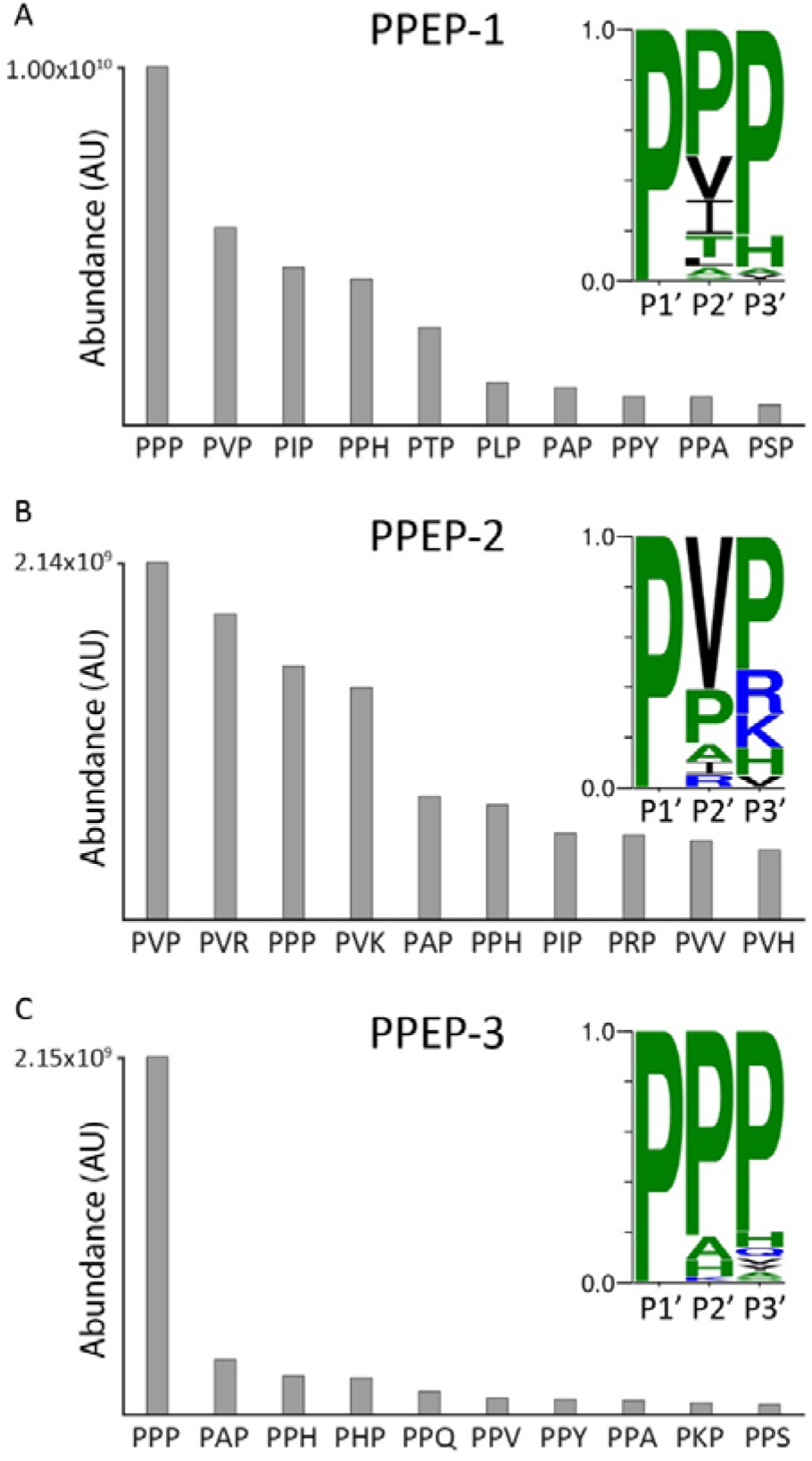
Top 10 most highly abundant 9-mer product peptides of PPEP-1, 2 and 3 reveal differences in prime-side specificity. The full combinatorial peptide library was incubated with recombinant PPEP-1, PPEP-2, or PPEP-3. Product peptides were analyzed using LC-MS/MS. Abundances were determined by summing the intensities of singly and doubly charged peptides. Discrimination between PXP and PPX peptides relied on both inspection of fragmentation spectra and C18 column separation (Supplementary Figs. 2 and 3). The 10 most highly abundant 9-mer product peptides formed by PPEP-1 (**a**), PPEP-2 (**b**) and PPEP-3 (**c**) and their abundances are represented as bars. A cleavage motif was constructed based on the relative intensities of the products peptides. The sequence on the X-axis represents the P1’-P3’ residues of the PXXGGLEEF product peptides.

The prime-side residues of the endogenous substrates of PPEP-1 (Fig. 1) were all represented among the top 10 product peptides, again demonstrating the feasibility of our method. In addition, the preference of PPEP-1 to hydrolyze substrates with three prolines at the P1’-P3’ (Fig. 3)^22^ was also demonstrated using the full combinatorial library (Fig. 4a). Interestingly, our approach revealed several previously unknown prime-side options that allow for cleavage by PPEP-1. The most striking findings included the cleavage of substrates that had either PPH, PPA, or PPY at their P1’-P3’ positions (Fig. 4a), since the presence of a Pro residue at P3’ was thought to be a determinant for proteolytic activity^20,22^. Therefore, it was initially hypothesized that the PHPGGLEEF/PPHGGLEEF product observed using MALDI-FT-ICR MS (Fig. 3) would in fact be PHPGGLEEF. However, manual inspection of the MS/MS fragmentation spectra revealed that PPEP-1 does tolerate PPH but not PHP at the P1’-P3’ sites. To corroborate this finding, we synthesized two FRET-quenched peptides (Lys_Dabcy_rEVNPPHPD-Glu_Edans_ and Lys_Dabcyl_-EVNPPPHD-Glu_Edans_), with either VNPPHP or VNPPPH as the core sequence, and tested these with PPEP-1. As expected, based on our library results, PPEP-1 is able to hydrolyze a VNP↓PPH, but not a VNP↓PHP peptide (Supplementary Fig. 4). Notwithstanding these exceptions, an overall preference of PPEP-1 for a Pro at the P3’ was observed (Fig. 4a).

For PPEP-2, substrate specificity had hitherto much less been explored. To a certain extent, PPEP-2 showed an overlapping specificity with PPEP-1 (Fig. 4b). For example, a high level of the PPPGGLEEF peptide was found and PPEP-2 also allows PPH at the P1’-P3’ positions. However, in line with the endogenous substrate (Fig. 1), PPEP-2 prefers a valine at the P2’ (Fig. 4b). Moreover, in contrast to PPEP-1, not all optimal substrates for PPEP-2 had at least two prolines at their P1’-P3’ positions. Of note, all peptides without prolines at the P2’ and P3’ positions had a Val at the P2’ position (Fig. 4b), again indicating that this is a strong determinant for PPEP-2 susceptibility (Fig. 1).

As mentioned above, we demonstrated for the first time that PPEP-3 is a genuine PPEP that cleaves Pro-Pro bonds (Fig. 4c). For PPEP-3, the most abundant product peptide corresponded to PPPGGLEEF (Fig. 4c). Since this peptide was relatively much more abundant than peptides with other amino acids at the P2’ and P3’ positions, this resulted in an overall motif that was dominated by proline at the P1’-P3’ positions. Still, PPEP-3 allowed several other residues at the P3’ that were not tolerated by the other two PPEPs. Furthermore, unlike the other PPEPs, PPEP-3 was able to cleave a PPHP motif (P1-P3’), as represented by the PHPGGLEEF product peptide (Fig. 4c).

Collectively, the above results showed that all three PPEPs preferred at least one proline at the P2’ or P3’ position. To emphasize the differences in such product peptides, extracted ion chromatograms (EIC) of every possible PXPGGLEEF/PPXGGLEEF peptide were constructed (Fig. 5). Not only does this clearly show the difference in product profiles, it also reveals the differences between PXP and PPX peptides such as PHP and PPH.

**Fig. 5.**
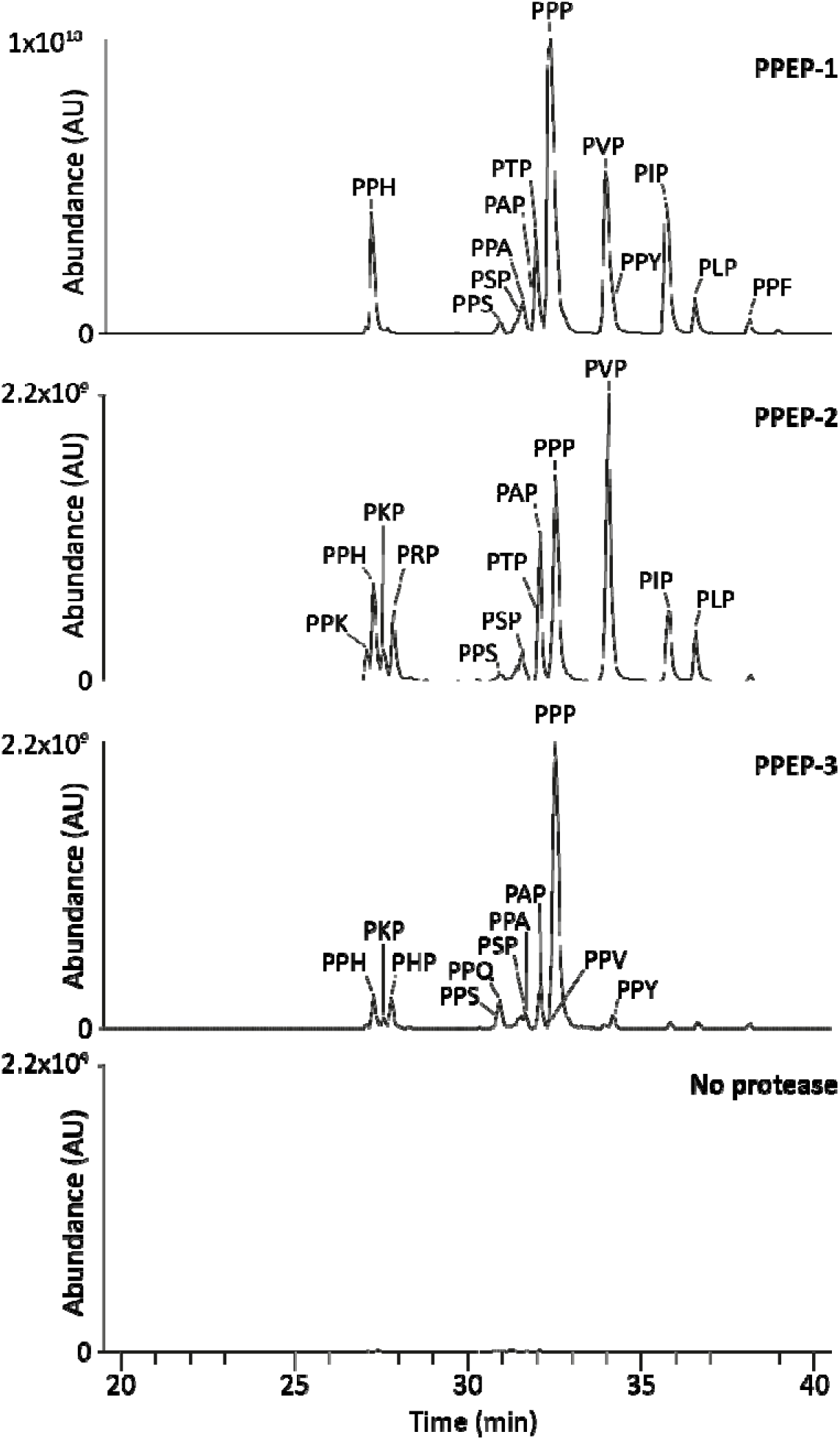
Extracted ion chromatograms of PXXGGLEEF product peptides after incubation with PPEPs reveal prime-side specificity profiles. The full combinatorial peptide library was incubated with each of the PPEPs for 3 h. A non-treated control was included to identify the amount of background peptides. After analysis of the product peptides using LC-MS/MS, EIC were constructed for all possible PXP/PPX product peptides (in total 19, both 1+ and 2+ *m/z* values were used). Discrimination between PXP and PPX peptides relied on both inspection of fragmentation spectra and separation on a C18 column (Supplementary Figs. 2 and 3). If product peptides were not separated on the column, lines indicate the relative abundances of the non-separated peptides. Mass tolerance was set to 10 ppm.

Based on the endogenous substrates (Fig. 1) and a small synthetic peptide library^22^, PPEP-1 was expected to only tolerate V, I, A and P at the P2’ position. To substantiate our results with the combinatorial peptide library, we synthesized twenty PPEP-1 FRET-quenched substrate peptides that only differed at the P2’ position (Lys_Dabcyl_EVNP↓PXPD-Glu_Edans_) and tested these with PPEP-1 in a time course kinetic assay. The results of these experiments are depicted in Fig. 6, in which substrates are ranked (from top left to bottom right) based on their increase in fluorescence during the 1 h incubation. Overall, these data (Fig. 6) correlated well with the results of the combinatorial library experiment (Fig. 5). Although cysteines were not included in the combinatorial library design (Fig. 2), the results with the VNPPCP FRET-peptide showed that it is not tolerated at the P2’ position by PPEP-1.

**Fig. 6.**
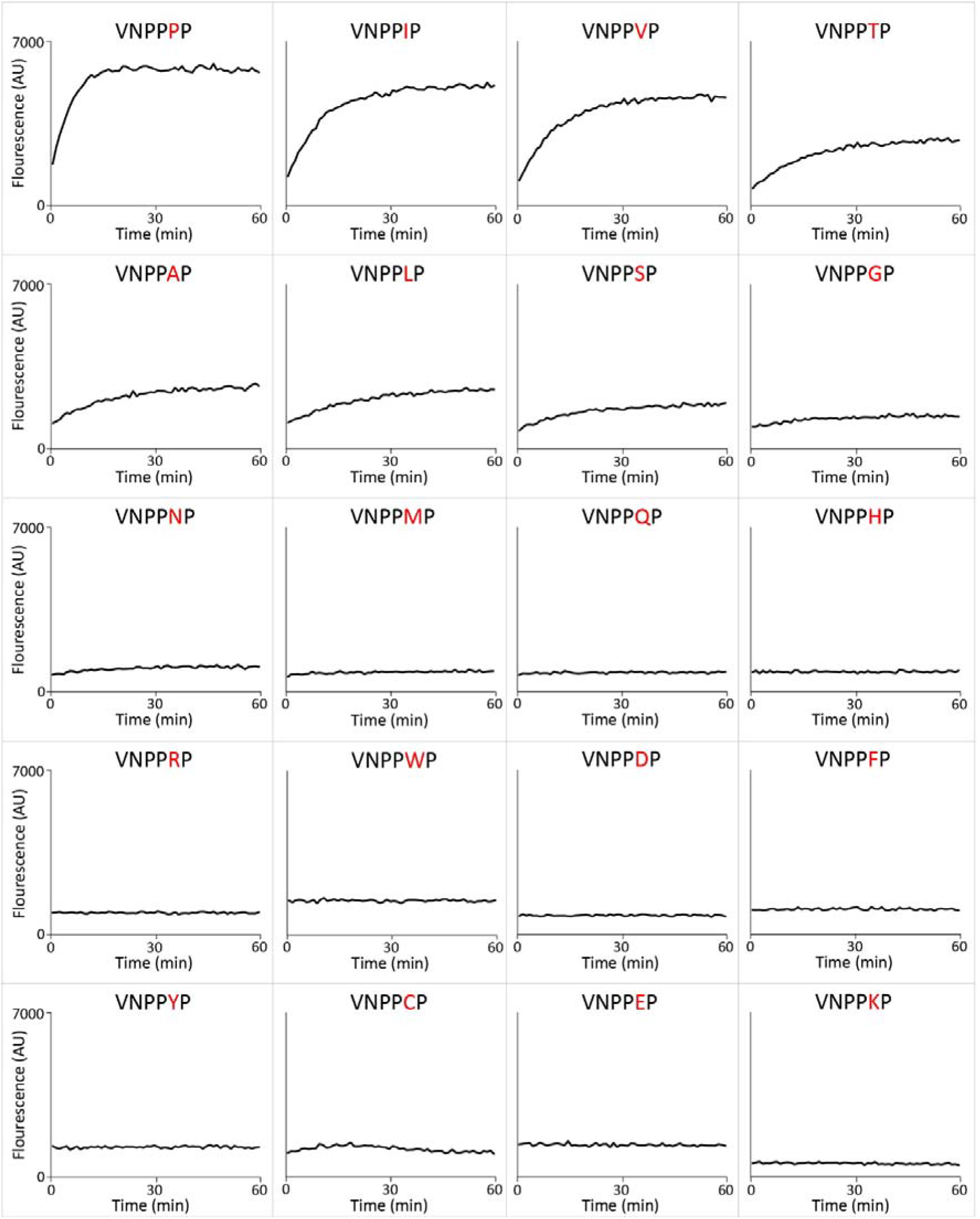
Time course of PPEP-1 mediated cleavage of synthetic FRET-quenched peptides with permutations at the P2’ position. The PPEP-1 substrate peptide VNP↓PVP was permutated to generate FRET-quenched peptides (Lys_Dabcyl_EVNPPXPD-Glu_Edans_) containing any of the standard 20 amino acids at the P2’ position. These peptides were incubated with PPEP-1 and fluorescence was measured during 1 hr. Peptides are sorted from top left to bottom right based on their cleavage efficiency.

### PPEP-3 is able to cleave endogenous PPEP-1 and PPEP-2 substrates when the valine at the P2’ position is replaced by a proline

The endogenous substrates of PPEP-1 and PPEP-2 contain the PVP motif at P1’-P3’ (Fig. 1) and the corresponding product peptides (PVPGGLEEF) were clearly observed using the combinatorial library approach (Fig. 5). However, this product peptide was not observed with PPEP-3 (Fig. 5), indicating that the corresponding PPEP-1 and PPEP-2 substrate peptides are most likely not cleaved by PPEP-3. We tested this hypothesis using two synthetic FRET-quenched substrate peptides, i.e. Lys_Dabcyl_EVNPPVPD-Glu_Edans_ and Lys_Dabcyl_-EPLPPVPD-Glu_Edans_, representing substrates of PPEP-1 and PPEP-2, respectively (Fig. 1). In line with our expectations, PPEP-3 did not hydrolyze either peptide (Fig. 7a). However, when the P2’ Val of both peptides was replaced by a Pro, cleavage by PPEP-3 did occur (Fig. 7a). On the contrary, although PPEP-1 and PPEP-2 can cleave peptides with four prolines at the P1-P3’position (Figs. 4a, b and 5), they can still not cleave each other’s substrate when the Val at the P2’ position is replaced by a proline (Fig. 7a).

**Fig. 7.**
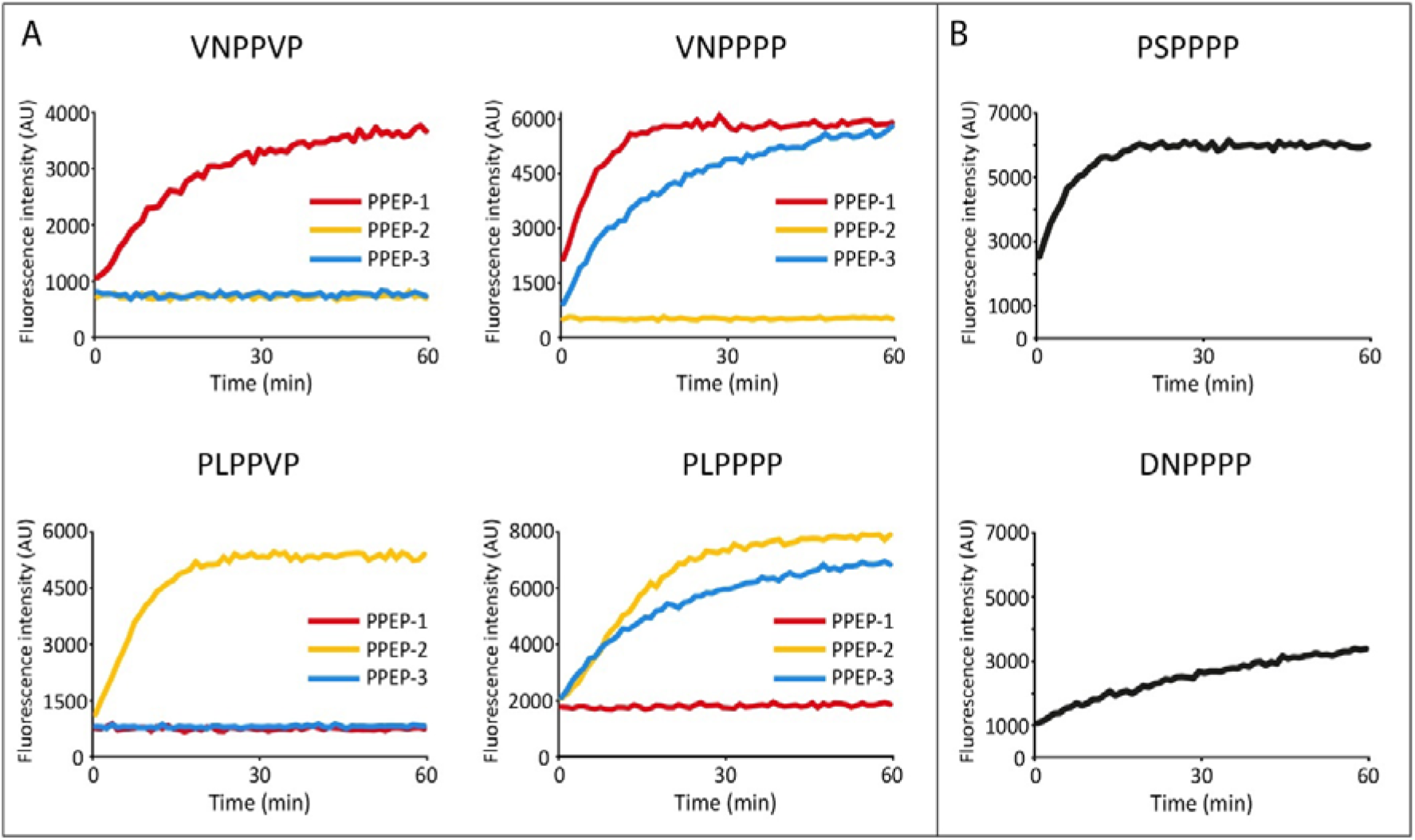
Time course of cleavage of synthetic FRET peptides by PPEP-1, PPEP-2 and PPEP-3. **a** Cleavage of PPEP-1 (Lys_Dabcyl_EVNP↓PVPD-Glu_Edans_) and PPEP-2 (Lys_Dabcyl_-EPLP↓PVPD-Glu_Edans_) substrate peptides, and their P2’=Pro variants, by PPEP-1, PPEP-2, and PPEP-3. **b** Cleavage of peptides containing cleavage motifs from putative *G. thermodenitrificans* PPEP-3 substrates by PPEP-3. Only the core sequences (P3-P3’) of the individual FRET-quenched peptides are indicated.

### Peptides with an XXPPPP motif as observed in *Geobacillus thermodenitrificans* proteins are cleaved by PPEP-3

Next, we looked for possible endogenous substrates of PPEP-3. For PPEP-1 and PPEP-2, genes encoding their substrates are found adjacent to the protease gene (Fig. 1). Next to PPEP-3, a gene encoding a protein (YpjP, Fig. 1) with three XXPPXX sequences is found (VTPPAS, EHPPQD and NTPPNW). In line with the data from the combinatorial library, corresponding FRET-quenched peptides were not cleaved by PPEP-3 (data not shown). Overall, our data from the library experiment indicate a strong preference of PPEP-3 for all prolines at the P1-P3’ positions (Fig. 4c, 5 and 7). Based on this observation, we hypothesized that possible endogenous substrates containing an XXP↓PPP motif are present in *G. thermodenitrificans* strain NG80-2. Indeed, *G. thermodenitrificans* encodes for four proteins containing four consecutive prolines, two of which contain a signal peptide for secretion as determined by DeepTMHMM and SignalP 6.0 (Supplementary Fig. 5)^27,28^. This last feature is thought to be of importance, since PPEP-3 itself is predicted to be a secreted protein. One of the identified proteins, GTNG_0956, contains both a putative CAP-domain as well as an SCP-domain. Admittedly, signal peptide prediction by SignalP 6.0 is inconclusive for this protein, since the signal peptide would be short in length and no cleavage site is predicted (Supplementary Fig. 5b). In contrast, DeepTMHMM predicts a signal peptide with higher confidence (Supplementary Fig. 5c). The other protein with an XXPPPP motif and a signal peptide is GTNG_3270. This protein is predicted with high confidence to possess a Sec/SPII signal sequence for integration in the lipid membrane. However, no functional domains were found for this protein. The putative PPEP-3 cleavage sites in GTNG_0956 and GTNG_3270 are PSP↓PPP and DNP↓PPP, respectively. We tested synthetic FRET-quenched peptides containing these motifs for cleavage by PPEP-3 (Fig. 7b). Both FRET peptides were indeed cleaved by PPEP-3, with PSPPPP being the optimal substrate of the two. Collectively, the above data show that the results from the library experiment resulted in testable hypotheses about possible endogenous PPEP-3 substrates in *G. thermodenitrificans* strain NG80-2.

## Discussion

A meticulous understanding of protease cleavage site specificity is important for the study and application of proteases in research, clinical, and industrial settings, e.g. when proteases are used as diagnostic biomarkers or for inhibitor design^5–7^. In addition to proteome wide approaches^29,30^, peptide libraries provide a means to explore protease specificity in detail. Usually this is done by testing large collections of possible peptide substrates^10,31^, for example as combinatorial peptide libraries^5,11^. One such method is similar to our method but uses Edman degradation instead of mass spectrometry to sequence the protease generated product peptides^32^. In addition to the lower sensitivity of this method, several amino acids could not be accurately detected and information on subsite cooperativity^33^ is lost. Our method takes each amino acid into account (with the exception of cysteine) and directly showed combinations of amino acids that were tolerated at the P2’ and P3’ positions. In the Positional Scanning Substrate Combinatorial Library (PS-SCL) approach, non-prime-sides are randomized while the proteolytic activity read-out depends on the release of a fluorophore, which is fixed and located at the P1’ position^5,11,18^. Since metalloprotease specificity, including that of PPEPs, is often also determined by prime-side residues^31,34^, such approaches are less applicable for this class of proteases. As opposed to our method, these methods allow for kinetic experiments. Although samples could be taken at different time points during the incubation, our method does not provide straightforward kinetic information. Still, our results showed clear preferences among the possible peptide substrates of PPEP-1, 2 and 3.

Others have been successful in profiling metalloprotease specificity using Proteomic Identification of Cleavage Site (PICS)^31,34^. Instead of using synthetic peptide libraries, this approach makes use of a peptide library derived from enzymatic digests of a proteome, for example human or *E. coli.* A major advantage of PICS is the ability to bioinformatically derive the non-prime-sides from the database searches. Although our experiments with PPEP-1 and the two P3-sublibraries showed that partial information about the non-prime side specificity can also be obtained with our method, we believe that a complementary XXPPXX library, in which the biotin is attached to the C-terminus of the peptides, is essential for a more comprehensive characterization of the non-prime-side specificity. Since the negative selection for substrates proceeds identically to that of the current library, both libraries can be mixed, allowing for the profiling of both the prime-side as well as the non-prime-side in a single experiment. In comparison with PICS, the major advantage of our method is that every possible substrate peptide is present in equimolar amounts. This feature enabled us to obtain an estimate of how well specific amino acids are tolerated at the prime-sides by comparing the relative abundances of the product peptides. However, the difference in intensities between the signals of the individual product peptides in the MS data also relate to how well these peptides are ionized, especially when extra basic amino acids are present, i.e. histidine, arginine and lysine^35^. We believe that this could explain the relative high contribution of these amino acids to the prime-side cleavage motifs that we have obtained.

In the Multiplex Substrate Profiling by Mass Spectrometry (MSP-MS) method^14^, a synthetic peptide library is used which was designed based on the idea that only two amino acids have to be positioned in an appropriate manner in a peptide substrate in order to be cleaved by a protease. Based on the inspection of the list of 228 peptides^36^, we predict that none of these would be cleaved by one of the PPEPs used in this study, making MSP-MS not suitable for specificity profiling of PPEPs.

The current method is straightforward in terms of sample preparation and LC-MS/MS analysis and based on a standard database search the activity of a PPEP could be readily determined. However, due to the high similarity between all peptides present in the library, we needed additional manual interpretation of the MS/MS data and testing of an additional small set of synthetic peptides in order to correctly assign several product peptides. Importantly, spectra of PXPGGLEEF peptides are dominated by the unique PGGLEEF (y_7_) fragment ion *(m/z* 748.351, Supplementary Fig. 2). This was for example essential in distinguishing PIPGGLEEF from PPIGGLEEF. The other unique fragment ion of PXPGGLEEF peptides, i.e. the b2 corresponding to PX, appeared less informative because it could also represent non-discriminatory internal fragments. We believe that this was one of the reasons why the results from the database searches were often ambiguous. Possibly other search algorithms, or training thereof, and new developments for prediction of tandem MS spectra^37^ could aid in the correct assignment of product peptides in terms of the amino acids at the second and third position in the PPEP-generated product peptides.

In addition to peptide fragmentation characteristics, separation of isomeric peptides using our reversed-phase chromatography system as part of the LC-MS/MS system was also essential. For example, we observed that peptides with an isoleucine elute earlier than the isomeric peptide having a leucine (Supplementary Figs. 3b, c), in line with what is known about the relative contribution of these two residues to the retention on a reversed phase column^38^. Moreover, PXP and PPX peptide pairs with an identical X residue that we have tested were well separated, with the exception of PIP and PPI (Fig. 5 and Supplementary Fig. 3). For example, histidine containing peptides were separated depending on the position of the histidine within the peptide, as also observed previously^39^.

Of the three PPEPs included in this study, PPEP-1 was so far the most extensively studied and we were able to confirm several key elements of its prime-side specificity^22,25^. However, we also had some unexpected findings. For example, when the P2’ subsite is occupied by a Pro, other residues than Pro are tolerated at the P3’, i.e. His, Phe and Tyr (Figs. 4a and 5). This was surprising, since we previously argued that a Pro at the P3’ position was a prerequisite for cleavage by PPEP-1^22^. The requirement for a Pro residue at P3’ was explained by the presence of a diverting loop in the co-crystal structure of PPEP-1 with a substrate peptide^20^. The Pro at P3’ aligns with Trp-103 of PPEP-1 due to a parallel aliphatic-aromatic interaction, thereby redirecting the remainder of the substrate (P4’ and onwards) out of the binding pocket by inducing a kink at the P2’ position. The ability of PPEP-1 to hydrolyze substrates with His, Phe, and Tyr at P3’ might be the result of similar aromatic-aromatic interactions (π-π stacking) with the Trp-103 and these residues^40^. Under this scenario, a Pro residue at the P2’ position is probably necessary to redirect the substrate from the diverting loop.

A bioinformatic analysis revealed multiple bacterial proteins with a domain predicted to have PPEP activity^23^. Remarkably, based on the amino acid residue at position 103 in PPEP-1, two groups were distinguished. In addition to PPEP-1, the Trp-103 group also includes PPEP-2. The other group, to which PPEP-3 belongs, has a Tyr at this position (Supplementary Fig. 6). Interestingly, a PPEP-1 W103Y mutant showed very low activity towards a substrate peptide as compared to WT^41^. For PPEP-2, the importance of this residue is less explored. Nevertheless, our data with PPEP-3 show that a tyrosine at this position is compatible with PPEP activity. Various product peptides were formed by PPEP-3, but their abundances were lower compared to those formed by the other two PPEPs. One explanation for this observation might be a more stringent non-prime-side specificity, since this would reduce the number of peptides with a preferable prime-side motif that can be cleaved. The importance of the non-prime side for PPEP-3 activity is supported by the FRET peptide cleavage assays, based on the two putative substrates (Fig. 7b). Alternatively, the difference in observed levels of product peptides in comparison to the other PPEPs might be due to a lower enzymatic activity of PPEP-3, for as yet unknown reasons. Whether the tyrosine in PPEP-3 that corresponds to the Trp-103 in PPEP-1 (Tyr-112, Supplementary Fig. 6) is responsible for the difference in prime-side specificity between PPEP-3 and the other two PPEP-s requires structural information, especially of a substrate-bound co-crystal.

Based on our data, PPEP-3 prefers all prolines at the P1-P4’ positions. Our search for this motif in the *G. thermodenitrificans* proteome resulted in the identification of two potentially secreted substrate proteins and two FRET-quenched peptides based on these potential substrates were indeed cleaved (Fig. 7b). Although no signal peptide cleavage site is predicted for GTNG_0956, the putative PPEP-3 cleavage site is located directly after the predicted signal peptide (Supplementary Fig. 5b). GTNG_3270 is predicted to be a lipoprotein and contains a putative PPEP-3 cleavage site close to the cysteine to which a diacylglyceryl moiety would be added for membrane anchoring (Supplementary Fig. 5b)^42^. For PPEP-1 and PPEP-2, the endogenous substrates were identified based on synthetic peptides, bio-informatic predictions and MS-based secretome analyses^21,25^. Interestingly, none of the sites in the endogenous substrates of these two PPEPs has four consecutive prolines, even though for both proteases the PPPGGLEEF product peptide was (one of) the major product peptides. In order to identify the endogenous substrate of PPEP-3, additional experiments such as secretome analyses in combination with gene knockout studies are needed, although we cannot exclude the possibility that the substrate(s) originates from a different organism than *G. thermodenitrificans*.

In conclusion, we present a method for protease specificity profiling which combines the complexity of a combinatorial peptide library with the high sensitivity of mass spectrometry analysis. We believe that the strategy presented here is a generic one which can, with a tailored design of the library, also be used to explore substrate specificities of other proteases. Importantly, with the new method we have not only confirmed the prime-side specificity of PPEP-1 and PPEP-2, but also revealed some new unexpected peptide substrates. Moreover, we have characterized a new PPEP (PPEP-3 from *Geobacillus thermodenitrificans*) which has a prime-side specificity that is very different from that of the other two PPEPs.

## Methods

### Expression and purification of PPEPs

PPEP-1 and PPEP-2 were expressed and purified as previously described^21,22^. For the expression of PPEP-3, a pET28a vector containing an *E. coli* codon optimized 6xHis-PPEP-3 (lacking the signal peptide) construct was ordered from Twist Bioscience. The pET-28a 6xHis-PPEP-3 plasmid was transformed to *E. coli* strain Rosetta and PPEP-3 expression was induced using 1 mM IPTG. Lysates were prepared as described in the protocol for preparation of cleared *E. coli* lysates under native conditions as described in the fifth edition of the QIAexpressionist (Qiagen). The lysates were loaded onto a 1 ml HisTrap HP column (GE healthcare) coupled to an ÄKTA Pure FPLC system (GE healthcare). Column was washed using wash buffer (50 mM NaH_2_PO_4_, 300 mM NaCl, 20 mM imidazole) and 6xHis-PPEP-3 was eluted using a step gradient with elution buffer (50 mM NaH_2_PO_4_, 300 mM NaCl, 500 mM imidazole). Imidazole was removed by dialysis using 50 mM NaH_2_PO_4_, 300 mM NaCl.

### Synthesis of the combinatorial peptide library

Combinatorial peptide libraries were synthesized basically as has been previously described^43^. In short, peptide libraries were synthesized by solid phase peptide synthesis on a Syro II peptide synthesizer (Multisyntech, Germany). Synthesis was performed in 19 reactors (2 ml) using about 1 g of Tentagel resin (Rapp-polymere, Germany) resin (total loading 190 μmol), applying Fmoc chemistry with HATU/NMM activation, 20 % piperidine in NMP for Fmoc removal and NMP as a solvent. For each fixed position in each reactor the same amino acid was coupled, for each random position (X) in each reactor a different amino acid was coupled, after which the resin beads were removed from each reactor, mixed thoroughly, and equally split over the 19 reactors again to allow for the subsequent stages of the synthesis. After the last random position, the resin beads were not mixed, leaving 19 sub-libraries. Cleavage using TFA/water/ethanethiol 18/1/1, 3h, RT, was used to isolate the peptides from the resin. Approx. 12 ml ether/pentane was added to each sublibrary and sub-libraries were incubated at −20 °C for 10 min before centrifugation at 3300 rpm for 10 min at −9 °C. Pellets were washed with approx. 13 ml ether/pentane and air-dried. Dried pellets were resuspended in 2 ml MilliQ/acetonitrile and freeze-dried. Stocks of 10 nmol peptide/μl were prepared in DMSO.

### Combinatorial peptide library assays

To remove non-biotinylated peptides, 50 nmol of peptides from the (sub)library (5 μl 10 nmol/μl stock in 1 ml PBS) was loaded onto a 3 ml filter column containing 1 ml Pierce Monomeric Avidin Agarose beads (Thermo) (binding capacity is >1.2 mg/ml biotinylated BSA or >18 nmol/ml). Prior to loading the libraries, the avidin column was washed five times with 1 ml 0.01% formic acid (pH 2.7) and subsequently washed five times with 1 ml PBS. After loading peptides, the flow-through was collected. Next, 1 ml PBS was loaded onto the column and flow-through was collected. Then, the collected flow-throughs were reapplied to the column to ensure saturation of the avidin beads. The column was washed five times with 1 ml PBS to remove non-biotinylated peptides. Next, 1 ml 0.1 M glycine (pH 2.7) was applied to the column and flow-through was discarded. Then, biotinylated peptides were eluted with 9 ml 0.1 M glycine (pH 2.7). Eluted peptides were desalted using reversed-phase solid phase extraction cartridges (Oasis HLB 1cc 30mg, Waters) and eluted with 400 μl 50% acetonitrile (v/v) in 0.1% formic acid. Samples were dried by vacuum concentration and stored at −20 °C until further use. If the binding efficiency of the avidin beads is the same for the peptide library as for biotinylated BSA, and no peptides are lost during the pre-wash steps, we expect approx. 20 nmol of peptide yield after the avidin pre-clearing step.

Pre-cleaned (sub)libraries (approx. 10 nmol) were incubated with a PPEP (200 ng) for 3 h at 37 °C in PBS. A non-treated control was included. After incubation, the samples were loaded onto an in-house constructed column consisting of a 200 μl pipette tip containing a filter and a packed column of 100 μl of Pierce High Capacity Streptavidin Agarose beads (Thermo, column was washed four times with 150 μl PBS prior to use), in order to remove the biotinylated peptides. The flow-through and 4 additional washes with 125 μL were collected. The resulting product peptides were desalted using reversed-phase solid phase extraction cartridges (Oasis HLB 1cc 30mg, Waters) and eluted with 400 μl 30% acetonitrile (v/v) in 0.1% formic acid. Samples were dried by vacuum concentration and stored at −20 °C until further use.

### MALDI-FT-ICR MS

To analyze samples using MALDI-FT-ICR MS, the vacuum dried product peptides were reconstituted in 20 μl 0.1% formic acid. Next, 1 μl sample was combined with 1 μl matrix (5 mg/ml α-Cyano-4-hydroxycinnamic acid) and 1 μl was spotted on an AnchorChip target (Bruker). Analysis was performed on a 15 T MALDI-FT-ICR MS (Bruker Daltonics).

### LC-MS/MS analyses

Product peptides were analyzed by on-line C18 nanoHPLC MS/MS with a system consisting of an Ultimate3000nano gradient HPLC system (Thermo, Bremen, Germany), and an Exploris480 mass spectrometer (Thermo). Fractions were injected onto a cartridge precolumn (300 μm × 5 mm, C18 PepMap, 5 μm, 100 A, and eluted via a homemade analytical nano-HPLC column (50 cm × 75 μm; Reprosil-Pur C18-AQ 1.9 μm, 120 A (Dr. Maisch, Ammerbuch, Germany). The gradient was run from 2% to 36% solvent B (20/80/0.1 water/acetonitrile/formic acid (FA) v/v) in 52 min. The nano-HPLC column was drawn to a tip of ~10 μm and acted as the electrospray needle of the MS source. The mass spectrometer was operated in data-dependent MS/MS mode for a cycle time of 3 seconds, with a HCD collision energy at 30 V and recording of the MS2 spectrum in the orbitrap, with a quadrupole isolation width of 1.2 Da. In the master scan (MS1) the resolution was 120,000, the scan range 350-1600, at standard AGC target @maximum fill time of 50 ms. A lock mass correction on the background ion m/z=445.12003 was used. Precursors were dynamically excluded after n=1 with an exclusion duration of 10 s, and with a precursor range of 10 ppm. Charge states 1-5 were included. For MS2 the first mass was set to 110 Da, and the MS2 scan resolution was 30,000 at an AGC target of 100% @maximum fill time of 60 ms.

### LC-MS/MS data analysis

We generated a database containing all 6859 peptides from the P3=Val sublibrary, i.e. Ahx-EVXPPXXGGLEEF. The Ahx in all peptide sequences was replaced by a Ile (they have an identical mass). Raw data were converted to peak lists using Proteome Discoverer version 2.4.0.305 (Thermo Electron), and submitted to the in-house created P3=Val sublibrary database using Mascot v. 2.2.7 (www.matrixscience.com) for peptide identification, using the Fixed Value PSM Validator. Mascot searches were with 5 ppm and 0.02 Da deviation for precursor and fragment mass, respectively, and no enzyme specificity was selected. Biotin on protein N-terminus was set as a variable modification. Raw data analysis was performed in Xcalibur Qual Browser (Thermo). The EIC displaying all PXPGGLEEF/PPXGGLEEF peptides was created by plotting the intensities of the signal corresponding to the monoisotopic *m/z* values of both 1+ and 2+ charged peptides. To assign individual peptides to their respective peaks, each individual peptide was plotted in an EIC and peptides were assigned to peaks based on retention time and abundance.

### FRET peptide cleavage assays

Time course kinetic experiments with PPEPs were performed using fluorescent FRET-quenched peptides. FRET peptides consisted of Lys_Dabcyl_EXXPPXXD-Glu_Edans_, in which X varied between the different peptides tested. To test cleavage of FRET peptides by PPEPs, 75 μL of FRET peptide (100 μM in PBS) was added to a well of a 96-well Cellstar black plate (Greiner). Immediately prior to the assay, 75 μl PBS containing a PPEP (0.2-1 μg) was added. Peptide cleavage was measured using the Envision 2105 Multimode Plate Reader. Fluorescence intensity was measured each minute for 1 h, with 10 flashes per measurement. The excitation and emission wavelengths were 350 nm and 510 nm, respectively. When comparing PPEP-1, PPEP-2, and PPEP-3 in a single experiment, the relative fluorescence was determined by regarding the highest signal as 100%.

### Bioinformatic analyses

PPEP-3 structure prediction was carried out using Colabfold^24^ with the following parameters: template mode=none, MSA mode=MMseqs2, pair mode=unpaired+paired, model type=auto, and number of recycles=3. Signal peptide predictions were performed using DeepTMHMM^27^ and SignalP 6.0^28^. For sequence alignments, the Clustal Omega Multiple Sequence Alignment tool was used^44^.

The cleavage motifs were created using Weblogo 3^45^ with the units set to probability. The sequences logos were generated based on the relative intensities of the 10 most abundant product peptides for each PPEP. A list with these 10 product peptides was created, in which each individual peptide occurred a number of times, according to its relative abundance to the other peptides. For example, if product ‘A’ was 100 times more abundant than product ‘B’, product ‘A’ was present 100 times more in this list than product ‘B’.

## Acknowledgements

This research was supported by an ENW-M grant (OCENW.KLEIN.103) from the Dutch Research Council (NWO). We thank prof. Ulrich Baumann for the critical reading of an earlier version of this manuscript. O.I.K. acknowledges support by the Interdisciplinary Scientific and Educational School of Moscow University ‘‘Molecular Technologies of the Living Systems and Synthetic Biology’’.

## Author contributions

P.J.H. and J.W.D. conceived the project. B.C., P.J.H. and J.C. performed experiments. B.C. and P.J.H. analyzed data. P.J.H., A.H.d.R. and A.O. performed mass spectrometry analyses. P.A.v.V. provided the means for mass spectrometry analyses. P.J.H., J.W.D. and R.A.C. designed the library. R.A.C. produced the library. J.C. and O.I.K. performed protein expression and purification. H.C.v.L. provided a protein expression construct. P.J.H. and P.A.v.V supervised the project. P.J.H. acquired funding. B.C. and P.J.H. visualized the results. B.C. and P.J.H. wrote the original draft. All authors reviewed and edited the paper.

## Competing interests

The authors declare no competing interests.

## Notes

### Competing Interest Statement

The authors have declared no competing interest.

