## Supplemental Figures for "In-depth specificity profiling of Pro-Pro endopeptidases (PPEPs) using combinatorial synthetic peptide libraries"

**Supplementary information**

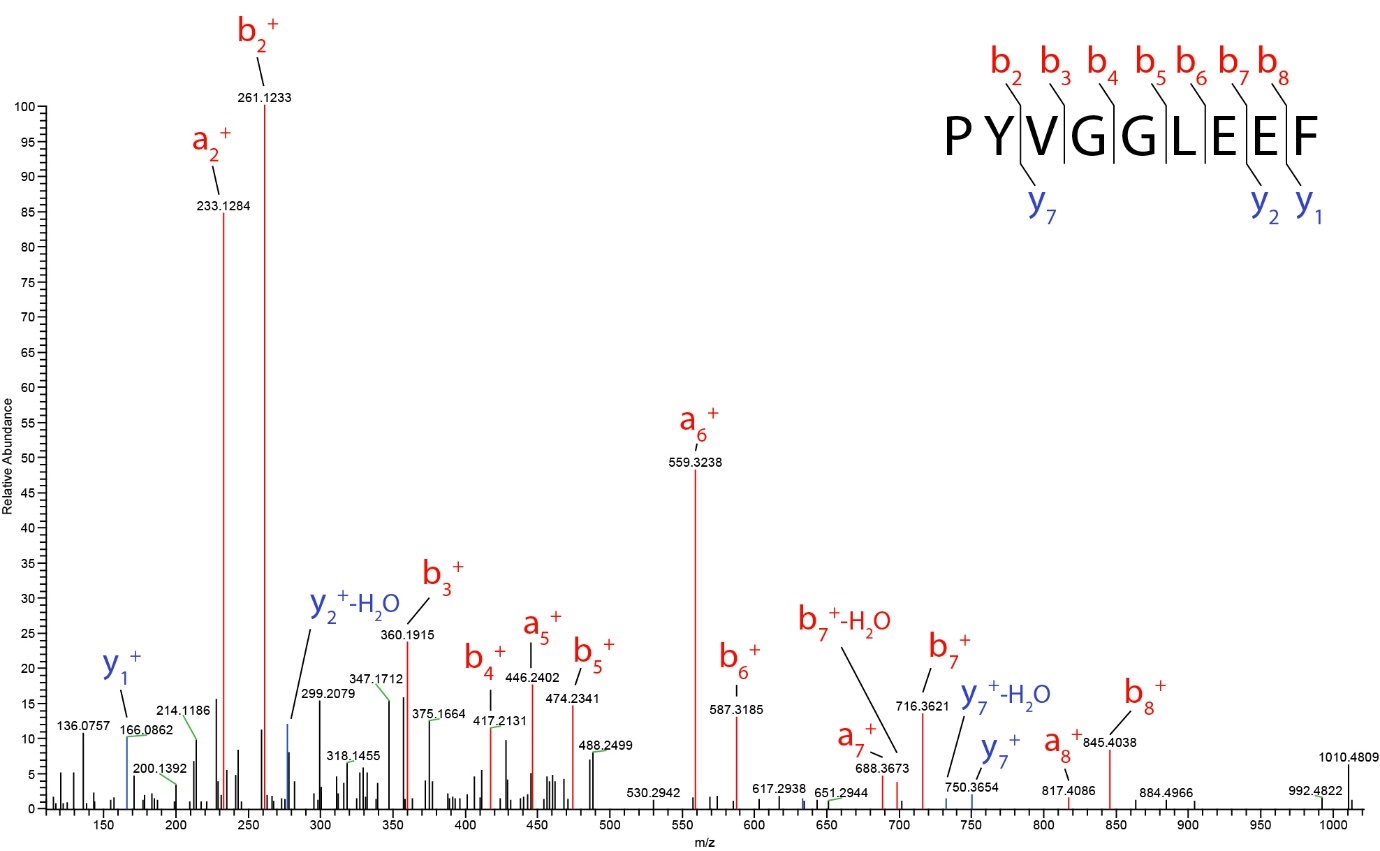

**Supplementary Fig. 1. Fragmentation spectrum of PYVGGLEEF***.* MS/MS spectrum of the PYVGGLEEF peptide that was used in the design of the combinatorial peptide library.

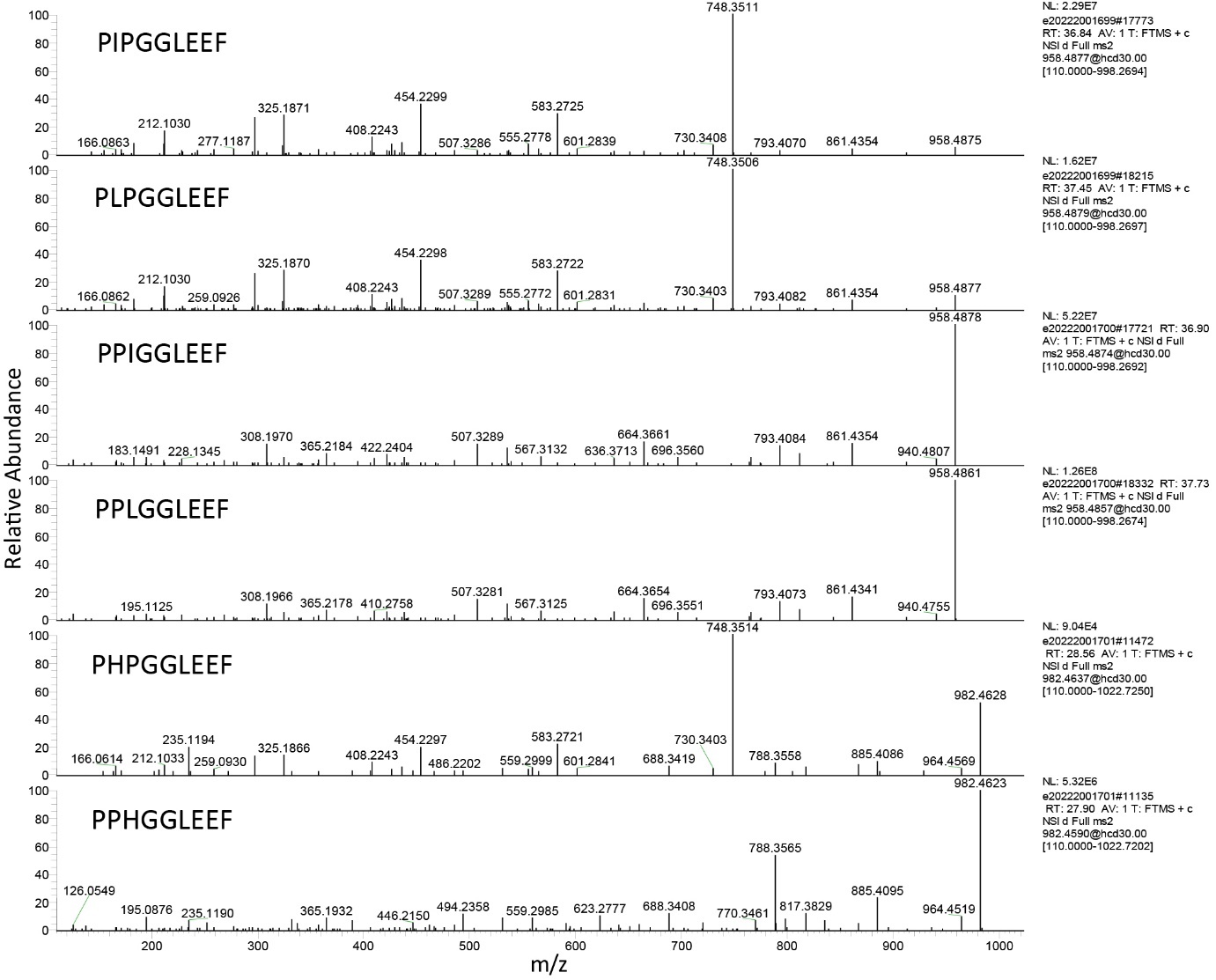

**Supplementary Fig. 2. Synthetic PXP/PPX product peptides display distinct fragmentation spectra.** Synthetic peptides with either a PXP or PPX motif were analyzed using LC-MS/MS.

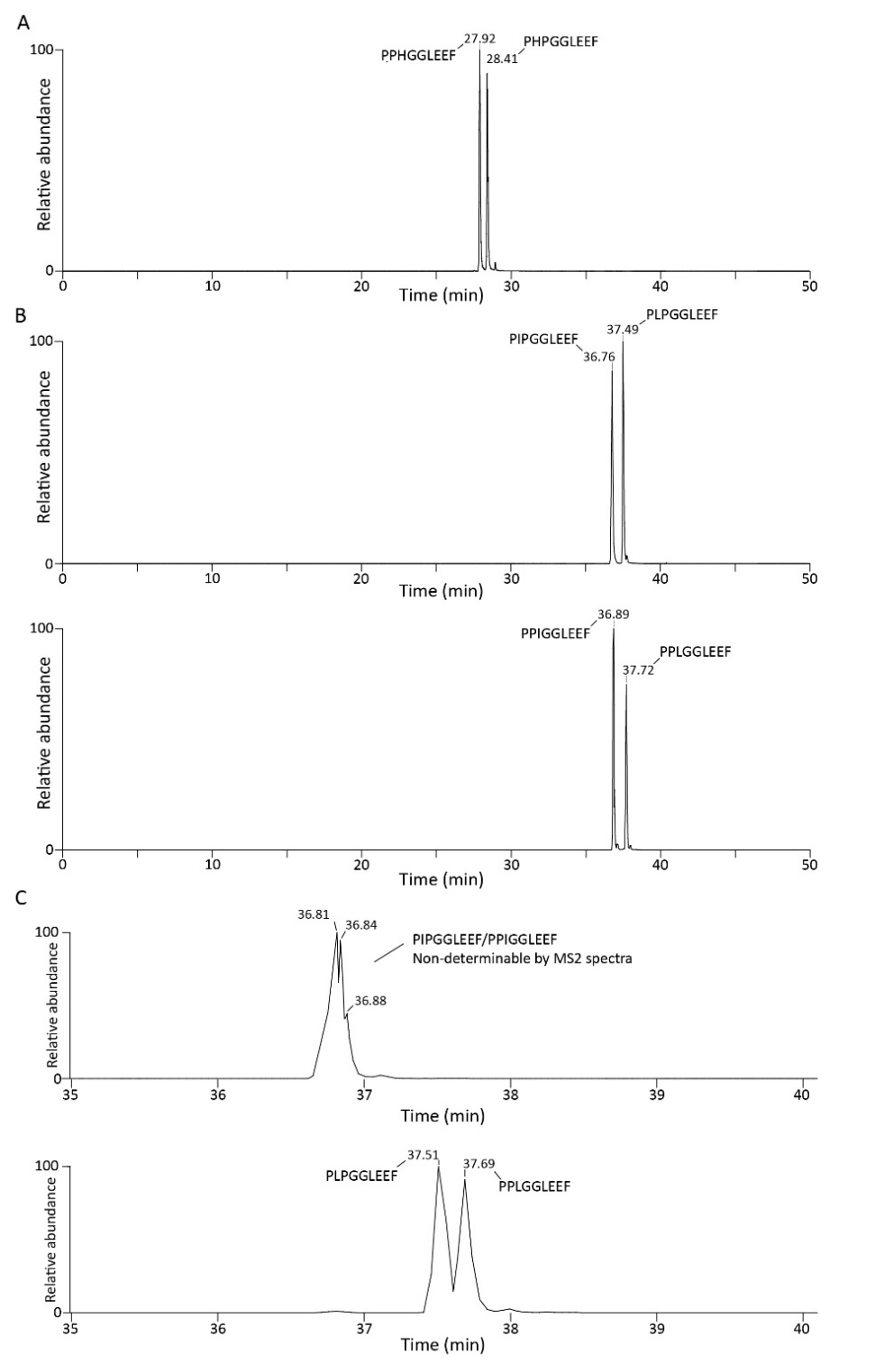
 **Supplementary Fig. 3. Separation of PXP/PPX peptides on C18 column.** Shown are extracted ion chromatograms of the corresponding peptides with a mass tolerance of 10 ppm. **A)** Synthetic peptides PHPGGLEEF and PPHGGLEEF were mixed at an equimolar concentration and analyzed using LC-MS/MS. Assignment of the peaks is based on LC-MS/MS analyses of the two peptides separately (data not shown). **B)** PXP/PPX peptides that contain either a Leu or Ile residue are separated on a C18 column. Assignment of the peaks is based on LC-MS/MS analyses of the two peptides separately (data not shown). **C)** PIPGGLEEF and PPIGGLEEF are not fully separated on a C18 column. Although some separation is observed, no conclusions could be made about the order of elution. However, panel B shows a lower retention time for PIPGGLEEF. PLPGGLEEF and PPLGGLEEF are, although not completely, separated on the column.

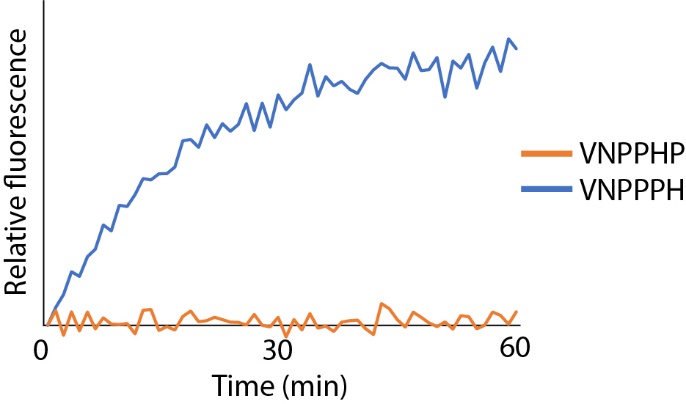

**Supplementary Fig. 4. PPEP-1 can cleave a VNPPPH peptide but not a VNPPHP peptide.** FRET-quenched peptides containing either VNPPHP or VNPPPH were incubated with PPEP-1 for 1 h. Fluorescence was measured in a fluorescence microplate reader.

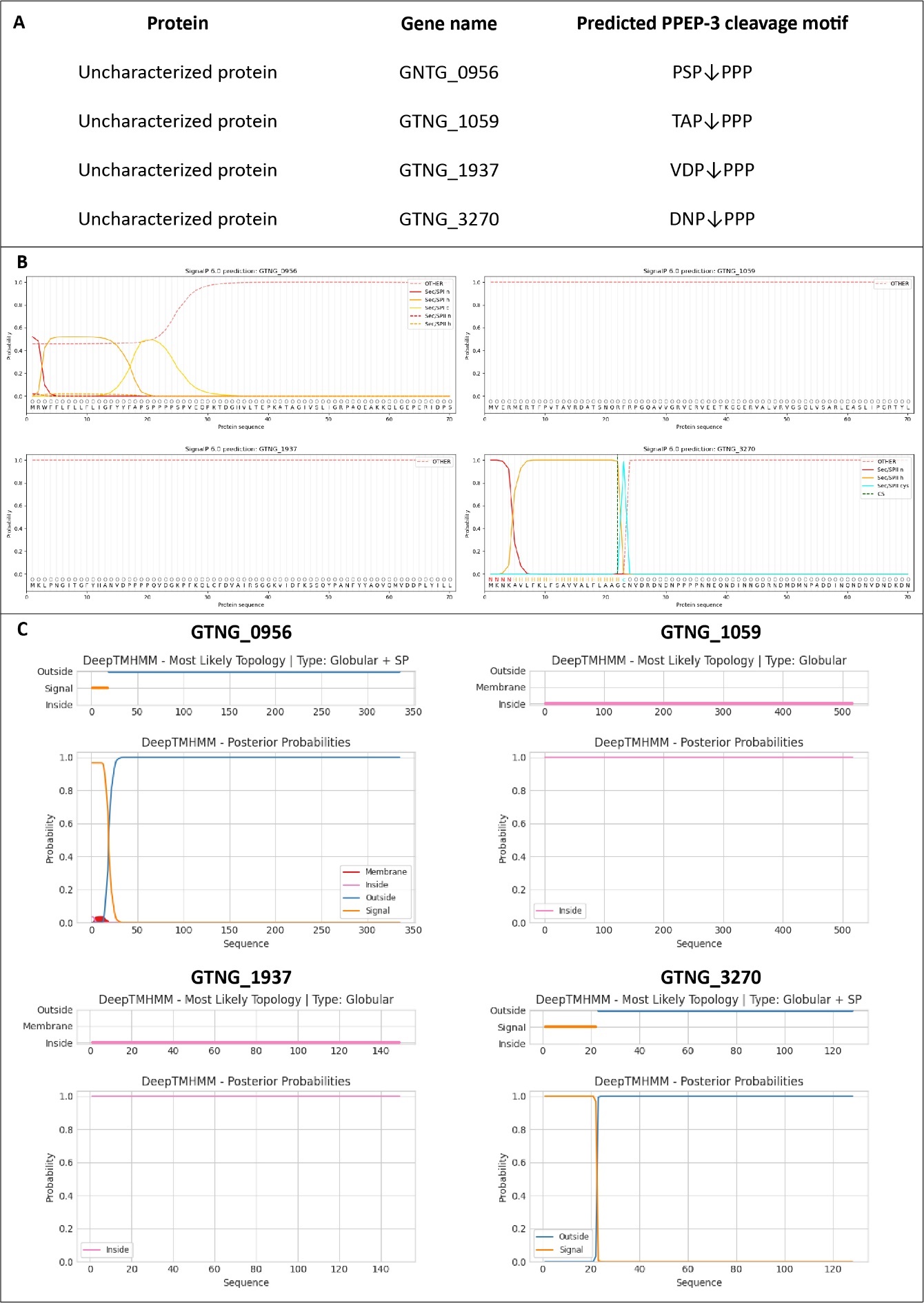
 **Supplementary Fig. 5. Signal peptide prediction of putative PPEP-3 substrates in G. thermodenitrificans.** **a** Overview of the proteins and their predicted PPEP-3 cleavage motifs. **b** Signal peptide prediction by SignalP 6.0. **c** Signal peptide prediction by DeepTMHMM.

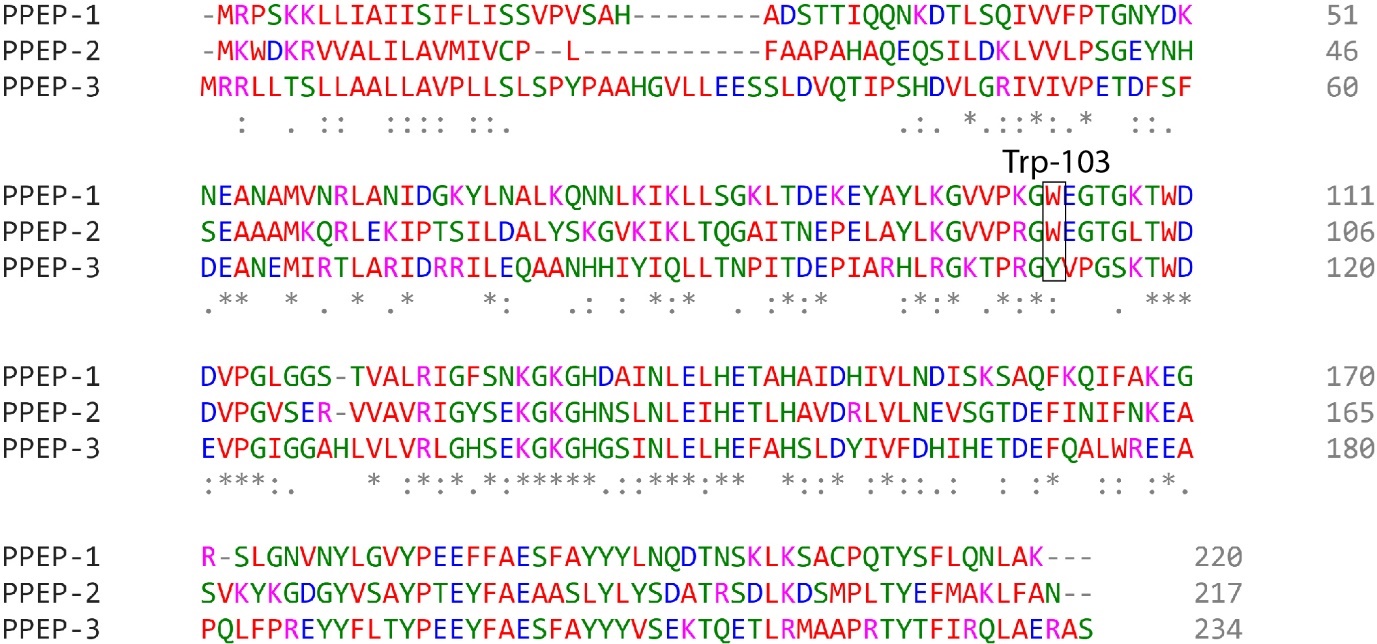

**Supplementary Fig. 6. Alignment of PPEP-1, PPEP-2 and PPEP-3.** Alignment was created using the Clustal Omega multiple sequence alignment tool.
